# Lecanemab Blocks the Effects of the Aβ/Fibrinogen Complex on Blood Clots and Synapse Toxicity in Organotypic Culture

**DOI:** 10.1101/2024.01.20.576458

**Authors:** Pradeep Kumar Singh, Elisa Nicoloso Simoes Pires, Zu-Lin Chen, Daniel Torrente, Marissa Calvano, Anurag Sharma, Sidney Strickland, Erin H. Norris

## Abstract

Proteinaceous brain inclusions, neuroinflammation, and vascular dysfunction are common pathologies in Alzheimer’s disease (AD). Vascular deficits include a compromised blood-brain barrier, which can lead to extravasation of blood proteins like fibrinogen into the brain. Fibrinogen’s interaction with the amyloid-beta (Aβ) peptide is known to worsen thrombotic and cerebrovascular pathways in AD. Lecanemab, an FDA-approved antibody therapy for AD, shows promising results in facilitating reduction of Aβ from the brain and slowing cognitive decline. Here we show that lecanemab blocks fibrinogen’s binding to Aβ protofibrils, normalizing Aβ/fibrinogen-mediated delayed fibrinolysis and clot abnormalities *in vitro* and in human plasma. Additionally, we show that lecanemab dissociates the Aβ/fibrinogen complex and prevents fibrinogen from exacerbating Aβ-induced synaptotoxicity in mouse organotypic hippocampal cultures. These findings reveal a possible protective mechanism by which lecanemab may slow disease progression in AD.

## Main

Alzheimer’s disease (AD) is a neurodegenerative dementia characterized by the accumulation of amyloid-beta (Aβ) aggregates in the brain parenchyma and in/around blood vessels (cerebral amyloid angiopathy, CAA)(1, 2). An early feature in AD is the disruption of the blood-brain barrier (BBB), which leads to the extravasation and accumulation of blood proteins within the brain, worsening AD pathology(1, 3, 4).

Fibrinogen, an abundant blood protein and major component of blood clots, can form a complex with Aβ upon binding to Aβ’s N-terminus (residues 8-20)(1, 5). Aβ may contribute to vascular abnormalities in AD by binding to fibrinogen and altering fibrin clot degradation(1). Consistent with this idea, reducing fibrinogen levels, or inhibiting Aβ/fibrinogen binding in AD mice leads to decreased BBB permeability, reduced neuroinflammation, decreased CAA, and less cognitive decline(6, 7).

The FDA-approved immunotherapy for AD, lecanemab, directed against Aβ protofibrils, reduces Aβ burden and slows cognitive decline in AD patients (8). However, little is known regarding its mechanisms-of-action. Lecanemab targets Aβ’s N-terminus (residues 1-16), overlapping with fibrinogen’s binding site on Aβ(2, 5).

We investigated the interaction between Aβ42 and fibrinogen in the absence and presence of lecanemab (Fig 1A-C). Lecanemab blocked Aβ42 binding to fibrinogen in a dose-dependent manner, while human IgG had no effect. The Aβ42 preparation used was comprised of curvy linear aggregates (small protofibrils) 30-90 nm in length (Fig 1D).

**Figure 1.**
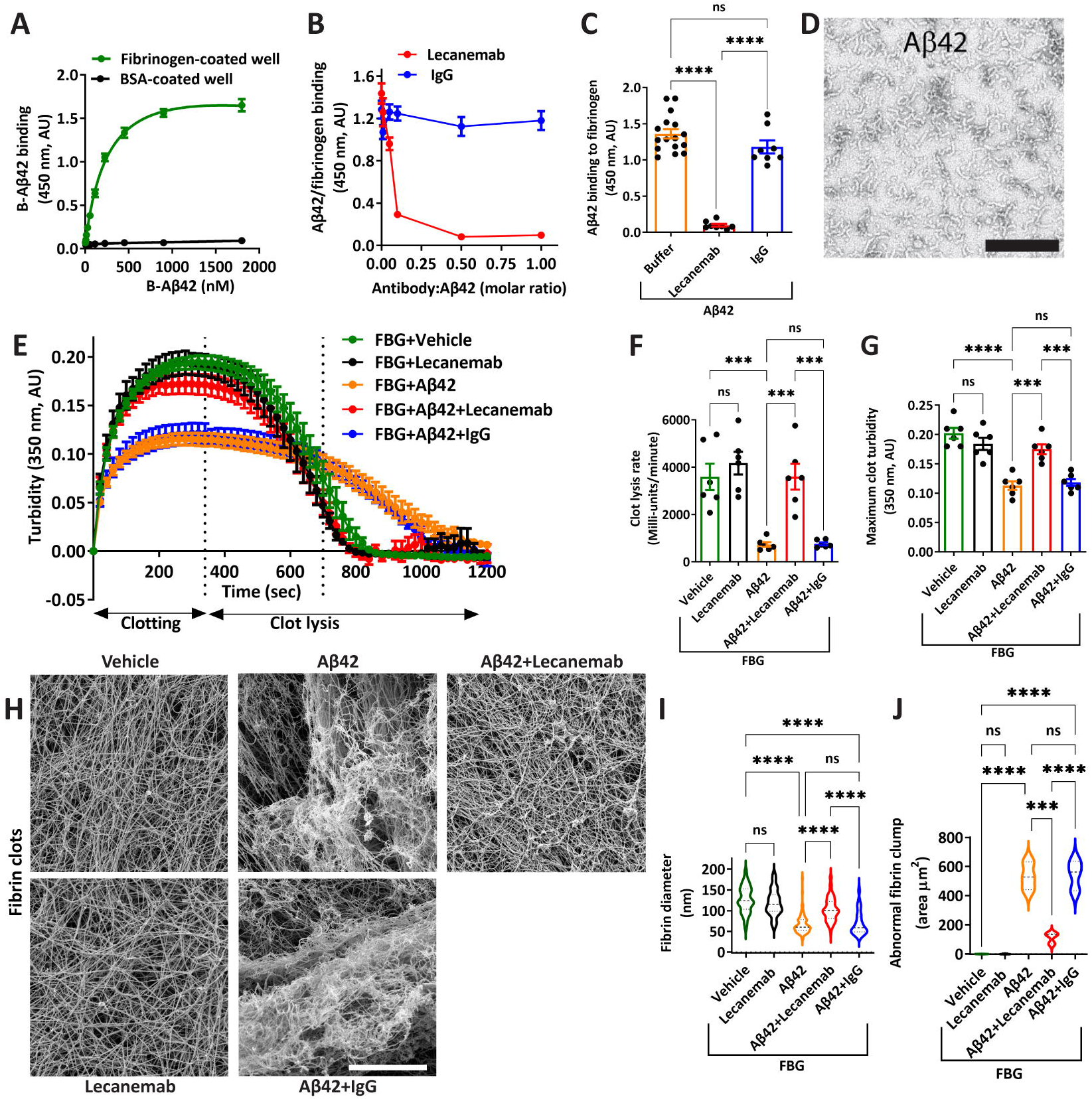
Lecanemab restores Aβ42 protofibril-induced delayed fibrinolysis and clot abnormalities by inhibiting the Aβ42/fibrinogen interaction *in vitro*. **(A)** Biotinylated Aβ42 protofibrils (B-Aβ42) bind to fibrinogen-coated wells. **(B)** Lecanemab dose-dependently blocked the binding of B-Aβ42 to fibrinogen, while human IgG control (IgG) had no effect. **(C)** Quantification of **B** (at equimolar ratio). **A-C:** Data from 3 independent experiments, n=8-16/group. **(D)** Transmission electron micrograph of Aβ42 protofibrils. Scale bar, 200 nm. **(E)** Turbidity assay shows clotting and clot lysis phases. Lecanemab, but not control IgG, corrected the Aβ42-induced delayed fibrinolysis. **(F)** Quantification of clot lysis rate in **E** (slopes between vertical dotted lines). **(G)** Quantification of maximum clot turbidity in **E. E-G:** Data from 6 independent experiments, n=6/group. **(H)** Representative scanning electron micrographs of fibrin clots from purified fibrinogen with different treatments. Scale bar, 10 μm. **(I, J)** Quantification of fibrin diameter and total area of abnormal fibrin clumps/clusters in clot images. Data from three independent experiments. Vehicle constitutes PBS+DMSO. Comparisons among multiple groups were performed using one-way ANOVA followed by Newman-Keuls multiple comparison test. Data are presented as mean ± SEM. ****p<0.0001, ***p<0.001; ns, not significant.

The interaction between Aβ42 and fibrinogen leads to an abnormal clot structure that is resistant to plasmin-induced fibrinolysis(1). Since lecanemab blocked the Aβ42/fibrinogen interaction, we analyzed the effect of lecanemab on Aβ42/fibrinogen-mediated impaired fibrinolysis in a purified protein system. As reported, Aβ42 protofibrils delayed plasmin-induced fibrinolysis. However, lecanemab blocked this effect of Aβ42, whereas human IgG did not (Fig 1E&F).

Aβ42 aggregates also alter fibrin clot turbidity, an indicator of altered fibrin assembly (7). Lecanemab, but not human IgG, rescued the defect in fibrin assembly caused by Aβ42 protofibrils (Fig 1E&G). We also analyzed clot morphology using scanning electron microscopy (SEM). As previously reported, Aβ42 disrupts normal clot morphology, causing thinning of the fibrin bundles, abnormal clustering, and entangled clumps (Fig 1H-J). However, in the presence of Aβ42 protofibrils and lecanemab, these structural clot abnormalities were significantly corrected (Fig 1H-J). Human IgG control had no effect on Aβ42-induced clot abnormalities (Fig 1H-J).

To determine if lecanemab inhibits the Aβ42/fibrinogen complex ex *vivo*, we incubated biotinylated Aβ42 protofibrils (B-Aβ42) with buffer, lecanemab, or human IgG and added them to normal human plasma (NHP). Immunoprecipitation was performed to pull down Aβ42 and any bound proteins. Fibrinogen immunoprecipitated with Aβ42 (Fig 2A&B). However, in the presence of lecanemab, Aβ42 did not pull-down fibrinogen, indicating that lecanemab blocked Aβ42/fibrinogen complex formation in NHP (Fig 2A&B). Also, consistent with the *in vitro* results (Fig 1E-J), lecanemab significantly corrected Aβ42-induced clot abnormalities in human plasma (Fig 2C-H).

**Figure 2.**
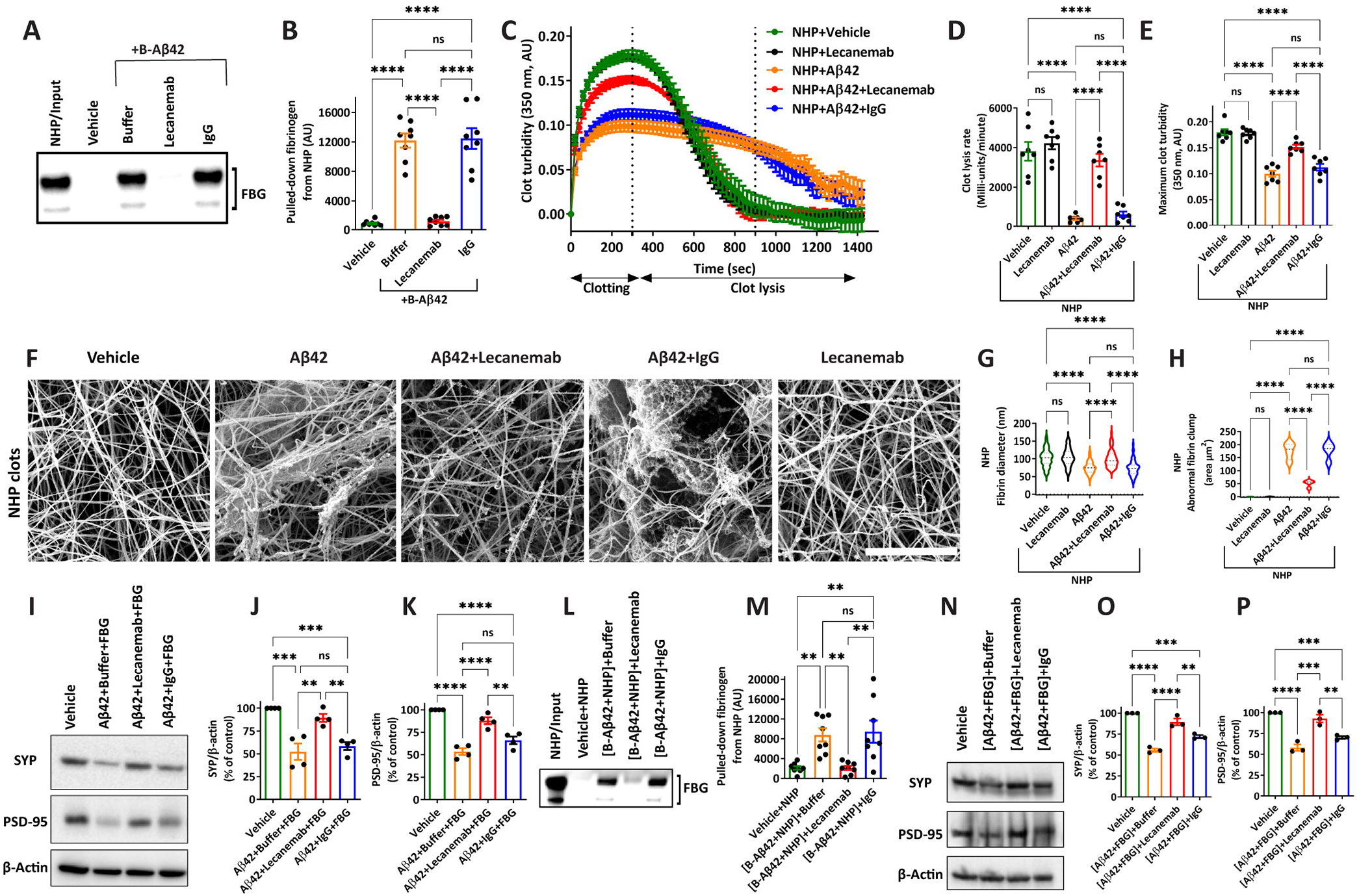
Lecanemab blocks Aβ42 protofibril-induced clot abnormalities and delays fibrinolysis in normal human plasma (NHP) and Aβ42/fibrinogen-mediated synaptotoxicity in mouse organotypic hippocampal culture (OHC). **(A)** Biotinylated Aβ protofibrils (B-Aβ42) immunoprecipitated fibrinogen from NHP but did not in the presence of lecanemab. Control IgG had no effect. **(B)** Quantification of **A**. Data from 3 independent experiments; n=8/group. **(C)** Clotting and fibrinolysis in NHP using turbidity assay. Lecanemab restored the Aβ42-induced delayed fibrinolysis in NHP. **(D)** Quantification of clot lysis rate in **C** (slopes between vertical dotted lines). **(E)** Quantification of maximum clot turbidity in **C. C-E:** Data from 7 experiments, n=7/group. **(F)** Representative scanning electron micrographs of clots formed from NHP with different treatments. Scale bar, 5 μm. **(G, H)** Analyses of scanning electron micrographs showing quantifications of fibrin diameter and total area of abnormal fibrin clumps/clusters. Data from 3 independent experiments. **(I)** Western blotting shows Aβ42+fibrinogen treatment reduced SYP and PSD-95 levels in OHC. However, in the presence of lecanemab, but not control IgG, the Aβ42/fibrinogen-mediated reduction in PSD-95 and SYP was minimized. **(J, K)** Quantification of SYP and PSD-95. Data from 4 experiments. Changes in synaptic markers were not due to cell death as determined by propidium iodide staining. **(L)** B-Aβ42 was added to NHP and incubated for one hour to form complexes with fibrinogen as shown in **A**. Lecanemab, buffer, or control IgG was then added. Western blot analysis of immunoprecipitation shows that lecanemab dissociated Aβ42/fibrinogen complexes. **(M)** Quantification of **L**. Data is from 3 independent experiments. **(N)** Western blotting shows preformed [Aβ42+fibrinogen] complex treatment reduced SYP and PSD-95 levels in OHC. However, treatment of preformed [Aβ42+fibrinogen] complex with lecanemab mitigated its synaptotoxicity. (**O, P)** Quantification of SYP and PSD-95. Data from 3 independent experiments. Vehicle constitutes PBS+DMSO. Comparisons among multiple groups were performed using one-way ANOVA followed by Newman-Keuls multiple comparison test. Data are presented as mean ± SEM. ****p<0.0001, ***p<0.001, **p<0.01; ns, not significant).

Synapse loss in AD is associated with memory impairment(4, 9). For example, the reduction of presynaptic protein synaptophysin (SYP) and post-synaptic density protein-95 (PSD-95) in the hippocampus corresponds to cognitive deficits in AD(10, 11). Extravasated fibrinogen can contribute to synaptic dysfunction(1, 3, 4, 12). Therefore, we explored if lecanemab could alter fibrinogen’s effect on Aβ42-mediated synaptotoxicity by examining the levels of SYP and PSD-95 in mouse organotypic hippocampal culture (OHC). Treatment of OHCs with a mixture of Aβ42 protofibrils and fibrinogen reduced SYP and PSD-95 (Fig 2I-K). Lecanemab, but not human IgG, inhibited Aβ42/fibrinogen-mediated synaptic changes (Fig 2I-K).

Lecanemab also dissociated preformed Aβ42/fibrinogen complexes in human plasma (Fig 2L&M). Moreover, lecanemab mitigated synaptotoxicity induced by preformed Aβ42/fibrinogen complexes in mouse OHCs (Fig 2N-P). Therefore, the ability of lecanemab to block Aβ/fibrinogen complex formation or dissociate the complex may be a component of its beneficial effects.

Amyloid-related imaging abnormalities (ARIA), a side effect of Aβ immunotherapy, are common in CAA(2, 13). Blocking the Aβ/fibrinogen interaction reduces CAA pathology and improves memory in AD mice(6). Our results show that lecanemab also targets the Aβ/fibrinogen complex. Although in some AD patients lecanemab treatment induces serious ARIA, it causes less ARIA than other anti-Aβ antibodies(2, 8, 14). Future studies are necessary to understand the connection between the Aβ/fibrinogen complex, lecanemab, and ARIA.

Our findings suggest that further investigations into lecanemab’s mechanisms-of-action are necessary in AD mouse models and AD patients. Studies include assessing lecanemab’s efficacy in dissociating and method of clearing Aβ/fibrinogen complexes *in vivo* and understanding its mechanism in mitigating Aβ/fibrinogen-induced synaptotoxicity. Moreover, given the neurodegenerative impact of extravasated fibrin(ogen) into the brain parenchyma independent of Aβ(1, 3, 4), exploring a combinatorial therapeutic strategy using lecanemab alongside a fibrin-specific antibody or another relevant target could be a promising treatment plan to improve upon the current AD immunotherapies (12).

## Materials and Methods

Details of reagents and methods (Aβ42 preparation, binding, immunoprecipitation, *in vitro* and *ex vivo* clotting and fibrinolysis, EM, and mouse OHCs) are included in the supporting information.

## Acknowledgements

This work was supported by NIH grant NS106668, Samuel Newhouse Foundation, John A. Herrmann, Zina Stern Fellowship, Mr. and Mrs. Pamela and Bill Michaelcheck, and May and Samuel Rudin Family Foundation.

