## Supplementary Information for "Lecanemab Blocks the Effects of the Aβ/Fibrinogen Complex on Blood Clots and Synapse Toxicity in Organotypic Culture"

**Author Contributions:** P.K.S, E.N.S.P., S.S., and E.H.N., designed research; P.K.S., E.N.S.P., Z.L.C., D.T., M.C., and A.S. performed experiments; P.K.S., S.S., and E.H.N., wrote the paper.

**Competing Interests:** The authors have no competing interests.

**Classification:** Biological Sciences/Neuroscience; NAS section: Biochemistry

**Keywords:** Alzheimer's disease, amyloid-beta, cerebral amyloid angiopathy, fibrinogen, fibrinolysis, lecanemab, vascular dysfunction

**This file includes:**

Supplementary Methods

### Materials and Methods

**Reagents and plasma-** The following reagents were used in these studies: Citrated normal human pooled plasma (NHP; George King Biomedicals, Inc); plasma-purified human fibrinogen (EMD Millipore); tissue plasminogen activator (alteplase; Genentech); human thrombin (Millipore-Sigma); Amyloid beta-42 peptide (Aβ42; Bachem and Anaspec); lecanemab (NDC 62856-212-01; Cardinal Health); human plasma-purified control IgG (Innovative Research). Plasminogen from normal human plasma was purified in-house (1).

**Preparation of Aβ42 protofibrils**- Aβ42 peptide was resuspended in ice-cold hexafluoroisopropanol (HFIP; Sigma Aldrich) and dried to form monomeric peptide films as described previously (2, 3). For Aβ42 soluble protofibril preparation, Aβ42 monomer films were dissolved in dimethyl sulfoxide (DMSO; Sigma), sonicated, and diluted in ice-cold PBS, pH 7.4. The final concentration of DMSO was 5% (v/v). The samples were incubated for 24 hours at $4^{o}$C in Protein LoBind tubes (Eppendorf). After incubation, the samples were centrifuged at 20,000xg for 10 minutes at $4^{o}$C to remove any insoluble Aβ aggregates. The protein concentration was determined using a Pierce BCA protein assay kit (Thermo Scientific)(2).

**Transmission electron microscopy (TEM)-** Morphology of soluble Aβ42 protofibril preparation was analyzed by transmission electron microscope (FEI, TECNAI G2) at Rockefeller University's Electron Microscopy Resource Center as described previously (2). TEM analysis showed that the Aβ42 preparation contained mostly elongated and curvy linear assemblies of 30-90 nm in length, which were previously defined as small protofibrils(4).

**Binding-** For ELISA-based binding, a 96-well plate was coated with human fibrinogen (250 ng/well) in binding buffer (0.1 M sodium bicarbonate buffer, pH 9.6). After washing, the plate was blocked using 1% BSA in PBS with 0.01% Tween-20 (PBST). Different concentrations of biotinylated Aβ42 (B-Aβ42) protofibrils were added, and the plate was incubated at $37^{o}$C for 15 minutes (for Fig 1A). To examine the effect of lecanemab or control IgG, B-Aβ42 (250 nM) was first incubated with buffer or different concentrations of antibodies before adding to a fibrinogen-coated 96-well plate (for Fig 1B). The plate was washed with PBST, and bound Aβ42 was detected using streptavidin-HRP. Three independent experiments were performed for this analysis.

**Immunoprecipitation-** Biotinylated Aβ42 protofibrils (B-Aβ42) were diluted in HBS and incubated with an equimolar concentration of antibodies (250 nM; lecanemab or control IgG). For reaction mixtures where antibodies were not added, PBS was included since the antibodies were prepared in PBS. Additionally, because B-Aβ42 was prepared in PBS containing 5% DMSO, an equivalent volume of PBS+DMSO (vehicle) was added to the reactions without B-Aβ42. After this step, normal human pooled plasma (NHP) was added to the reaction mixture and incubated for an additional hour at room temperature. Dynabeads M-280 Streptavidin (Invitrogen) were used to pull down B-Aβ42 and its bound proteins according to the manufacturer's instructions. The level of fibrinogen pulled down with different treatments was analyzed by Western blotting (under reducing conditions) using an anti-fibrinogen antibody (Dako). Three independent experiments were performed for these analyses.

To prepare Aβ42/fibrinogen complexes in NHP and then dissociate them using lecanemab, B-Aβ42 or vehicle (PBS+DMSO) was initially added to NHP diluted in HBS and incubated for 1 hour at room temperature. Then PBS or antibody (lecanemab, control IgG) was added to the reaction mixture and incubated for 2 hours at room temperature. B-Aβ42 and antibodies were in equimolar ratio. Dynabeads M-280 Streptavidin (Invitrogen) were used to pull down the Aβ42/fibrinogen complex, which was then analyzed using Western blotting as described above. Three independent experiments were performed for these analyses.

***In vitro* clotting and fibrinolysis-** Fibrinogen stock solution was prepared in HEPES buffered saline-HBS (20 mM HEPES, pH 7.4, 140 mM NaCl) as described previously (5). For clot formation and lysis, Aβ42 protofibrils were first incubated with PBS, lecanemab, or control human IgG in HBS for 15 minutes at 37$^{o}$C in Protein LoBind tubes (Eppendorf) on a nutator. PBS was included since the antibodies were prepared in PBS. Reaction mixtures without Aβ42 protofibrils contained either vehicle (PBS+DMSO) or vehicle+lecanemab in HBS. After incubation, fibrinogen, tissue plasminogen activator (tPA), and plasminogen were added to these reaction mixtures and incubated for an additional 15 minutes. The clotting was initiated in a 96-well plate by adding thrombin and calcium chloride to incubated reaction mixture (1, 2). Thrombin converts fibrinogen into fibrin, increasing turbidity (clotting phase). Maximum turbidity marks the completion of clotting. After the clotting phase, tPA converts plasminogen into plasmin, initiating fibrin clot degradation and decreasing turbidity (clot lysis/fibrinolysis phase) (1). Clot formation and dissolution were monitored by measuring the changes in turbidity (optical density) at 350 nm over time using a spectrophotometer (Molecular Devices) at 37$^{o}$C (1). The clot lysis rate (change in optical density or transmittance in the lysis phase) was calculated for each treatment. The clot lysis rate was calculated using the slopes between 340-700 seconds of curves (fibrinolysis phase) using SoftMax Pro 7.1 software (Molecular Devices). In the reaction mixture, the final concentration of Aβ42 and antibodies was 3 µM, fibrinogen was 1.5 µM, plasminogen was 150 nM, tPA was 750 pM, thrombin was 1.5 U/ml, and calcium chloride was 3.75 mM. Six independent experiments were performed for this analysis.

***Ex vivo* clotting and fibrinolysis in NHP-** For clot formation and lysis, Aβ42 protofibrils were first incubated with PBS, lecanemab, or control human IgG in HEPES buffered saline-HBS (20 mM HEPES, pH 7.4, 140 mM NaCl) for 15 minutes at 37$^{o}$C in Protein LoBind tubes (Eppendorf) on a nutator. Reaction mixtures without Aβ42 protofibrils contained either vehicle (PBS+DMSO) or vehicle+lecanemab in HBS. After incubation, NHP and tPA were added to these reaction mixtures and incubated for an additional 15 minutes. Clotting was initiated in a 96-well plate by adding thrombin and calcium chloride. Clot formation and dissolution were monitored using a spectrophotometer (Molecular Devices) as described above. The clot lysis rate was calculated using the slopes between 300-900 seconds of curves (fibrinolysis phase) as described above. In the reaction mixture, the final concentration of Aβ42 or antibody was 3 µM, fibrinogen was 1.5 µM, exogenous tPA was 750 pM, and thrombin was 1.5 IU/ml. The final dilution of NHP was 6-fold in the reaction mixture (16.6 µl in 100 µl final volume). Seven independent experiments were performed for this analysis.

**Scanning electron microscopy (SEM)-** Aβ42 protofibrils were first incubated with PBS, lecanemab, or control human IgG in HEPES-buffered saline (HBS; 20 mM HEPES, pH 7.4, 140 mM NaCl) for 15 minutes at 37$^{o}$C in Protein LoBind tubes (Eppendorf) on a nutator. Reaction mixtures without Aβ42 protofibrils contained either vehicle (PBS+DMSO) or vehicle+lecanemab in HBS. After incubation, purified fibrinogen was added. For clot analysis in human plasma *ex vivo*, NHP was added instead of fibrinogen. The samples were incubated for an additional 15 minutes. For SEM, clots were prepared on round glass coverslips at room temperature. The incubated clotting reaction mixtures with different treatments (70 µl) were gently spotted on clean coverslips. Thrombin and calcium chloride (30 µl) was gently added. The final dilution of NHP was 6-fold in the reaction mixture (16.6 µl in 100 µl final volume). In the purified system, the final concentration of fibrinogen was 1.5 µM. The final concentration of Aβ42 and antibodies was 3 µM, thrombin was 1.5 IU/ml, and calcium chloride was 3.75 mM. After 1 hour, the clots were washed with sodium cacodylate buffer (0.1M, pH 7.2) and fixed with 2% glutaraldehyde (in sodium cacodylate buffer) for 30 minutes. Clots were washed after fixation, serially dehydrated with graded series of ice-cold ethanol (30%, 50%, 70%, 90%, 100%), critical point dried, and sputter coated as described previously (2, 5). Imaging was done using a scanning electron microscope (JSM-IT500HR, JEOL) at Rockefeller's Electron Microscopy Resource Center. Images from three different experiments were used to quantify diameter of fibrin strands and total area of abnormal fibrin clumps using Image J software as described previously (1).

**Organotypic hippocampal culture (OHC)-** OHCs were prepared according to Stoppini et al(6). with slight modifications (7). Briefly, post-natal day 8-10 C57BL/6 mice were decapitated instantaneously, and their brains were quickly removed from the skull and washed with ice-cold HBSS. The hippocampi were dissected, sliced into 400 µm-thick slices, and arranged on organotypic inserts in six-well cell culture plates. Each insert contained two hippocampal slices from three different animals (six slices total). The slices were cultured in an incubator at 37°C with 5% $CO_2$ using an interface method. The culture medium consisted of a mixture of minimum essential medium, HBSS, heat-inactivated horse serum, D-glucose, HEPES, NaHCO3, and antibiotics. Medium changes were performed every 3-4 days. The slices were maintained for 14 days *in vitro* before experimentation. After 14 days, hippocampal slice cultures were exposed for 24 hours at 37°C and 5% $CO_2$ with different treatments.

For treatment, Aβ42 protofibrils were added in culture media and first incubated with PBS, lecanemab, or control IgG for 1 hour at 37°C. Culture media without Aβ42 protofibrils was incubated in the same way with vehicle (PBS+DMSO). These preparations were then mixed with fibrinogen or buffer and further incubated for 1 hour at 37°C before adding them to culture. The final concentration of fibrinogen was 50 nM. The final concentration of Aβ42 or antibody (lecanemab or IgG) was 150 nM.

To assess the effect of lecanemab on preformed Aβ42/fibrinogen complexes, Aβ42 protofibrils were first incubated with fibrinogen for 1 hour at 37°C, followed by the addition of antibody and further incubation for 1 hour at 37°C. These reaction mixtures were used to treat the hippocampal slice cultures. The final concentration of Aβ42 protofibrils or antibody (lecanemab or IgG) was 150 nM. At the end of treatment (24 hours), slices were rinsed with PBS, lysed in ice-cold RIPA buffer containing 25 mM Tris-HCl, pH 7.6, 150 mM NaCl, 1% NP-40, 1% sodium deoxycholate, 0.1% SDS, and protease and phosphatase inhibitors (Thermo Scientific) and homogenized by vigorous vortex shaking. Protein concentration was estimated by Pierce BCA protein assay kit (Thermo Scientific) (2). Homogenate samples (30 µg) were loaded onto separate lanes of SDS-PAGE gels and transferred to PVDF membrane for Western blotting. The membranes were probed for synaptophysin (SYN) and post-synaptic density protein-95 (PSD-95) using anti-synaptophysin and anti-PSD-95 antibodies (Abcam), respectively. Anti-β-actin antibody (Sigma) was used to probe β-actin level as a normalization control.

Cell degeneration was assessed with a slight modification as previously described (7). Twenty-three hours after treatment, 5 µM propidium iodide (PI) was added to the culture medium and incubated for 1 hour in an incubator at 37°C with 5% $CO_2$. Fluorescence imaging of the uptake of PI integrated over the whole slice was examined on EchoRevolve microscope (Echo, San Diego, CA) and analyzed using NIH ImageJ software.

**Statistical analyses-** All statistical analyses were performed using GraphPad Prism 9 software. Comparisons among multiple groups were performed using one-way ANOVA followed by Newman-Keuls multiple comparison test. (****$p < 0.0001$, ***$p < 0.001$, **$p < 0.01$, ns, not significant). Results were obtained from several independent experiments ($n \geq 3$). Samples were randomized and analyses were blinded. There were no samples excluded from the analysis.
